# A Whole-Genome Association Approach for Large-scaled Inter-species Trait

**DOI:** 10.1101/454363

**Authors:** Qi Wu, Huizhong Fan, Lei Chen, Yibo Hu, Fuwen Wei

## Abstract

Genome wide association studies (GWAS) have provided an avenue for the association between common genetic variants and complex traits. However, using SNP as a genetic marker, GWAS has been confined to detect genetic basis traits only for within species but not for the large-scale inter-species traits. Here, we propose a practical statistical approach that is using kmer frequencies as the genetic markers to associate genetic variants with large scale inter-species traits. We applied this new approach to the trait of chromosome number in 96 mammalian proteomes, and we prioritized 130 genes including *TP53* and *BAD*, of which 6 were candidate genes. These genes were proved to be associated with cellular reaction of DNA double-strand breaks caused by chromosome fission/fusion. Our study provides a new effective genomic strategy to perform association studies for large-scaled inter-species traits, using the chromosome number as a case. We hope this approach could provide exploration for broadly widely traits.

## Introduction

With the development of sequencing technologies, GWAS have been proven powerful and increasingly affordable tool for scaning and detecting candidate genes responsible for complex disease in human or important economically traits in plants or domestic animals [1-4]. Generally speaking, it is the central aim of genetics that to understand the association between heritable variations and phenotypic changes [5]. At present, all of the GWAS are focused on the genetic variants in hundreds or thousands of individuals under species. It is an interesting question to ask that whether GWAS is available between genetic variants and large-scale inter-species traits. In practice, the major problem is lack of an approach to bridge the genome data to the heritable traits. This entails two close related issues: firstly, a marker is needed to quantify various difference of genome/proteome sequences among species. Secondly, a statistical method is needed to assess the association between the makers and the large-scale inter-species traits.

For the first issue, we are able to identify the probable maker to represent the characteristics of different genomes with the help of alignment-free (AF) sequence comparison analysis [6-9]. Broadly speaking, AF methods focuses on the subsequences (words or kmer) of defined or varied length from the sequence, which their types [6], frequencies [7], or position [10] of the kmers on the sequence were considered as characteristics. Various compositional heterogeneity of sequence, including CpG islands, transposon elements, centromere as well as telomere in the genome could be delicately reflected in the characteristics of kmers [11]. Moreover, all types of variations such as substitution, insertion/deletion, duplication as well as rearrangements could also be recorded in the characteristic of subsequences [12]. For quantifying the overall feature of the sequence, AF methods compares and analyzes the characteristics between the concerned sequences. In practice, many excellent AF sequence comparison studies have been successfully performed to solve the problems of whole-genome phylogeny construction. The results of AF comparison studies are in correspond with those using morphological data or molecular data [7,9,12,13]. Therefore, it is reasonable to deduce that the kmer characteristics of the genome can describe the sequence variations which determine the large-scaled inter-species traits. This suggests the possibility of using kmer characteristics as the genetic markers to preform GWAS analysis on inter-species traits.

Numerous strategies and statistical approaches have been developed to meet the technical challenges and the great opportunities provided by GWAS [4,14,15]. One of such approaches is gene-based GWAS analysis [14]. In GWAS analysis, gene-based association analysis could combine the significance of all SNPs in a gene and effectively assess the association between gene and phenotype traits in hundreds or thousands of individuals. As such, it is possible to combine the significance of all kmer frequencies in a gene and assess the significance of the association between the gene and the inter-species traits in the evolutionary studies. Thus, the candidate genes that have significant effects on the large-scale inter-species traits during the evolution could be identified. Moreover, by mapping the significant kmers to the gene sequences, the potential functional domain regions in the genes associated with the traits could also be identified.

In this study, we proposed a practical statistical approach that use kmer frequency as the genetic markers to quantify the overall statistical feature of the genomes/proteomes among different species, and to associate genetic variants with large scale inter-species traits. The proteome data of 96 mammals was used as samples. The chromosome number of these species and the kmer frequencies of each proteome were used as the phenotypic variables and the genetic markers respectively. A total of 130 genes including *TP53* and *BAD* were identified significantly associated with the chromosome variations. The annotation results of these genes show that they are related to the cellular reaction of DNA double-strand breaks caused by chromosome fission/fusion. Our study provides a new approach to perform GWAS on the large-scale inter-species trait, and it was applied to analysis the genetic variants that are associated with chromosome number during the evolution of species. In fact, besides chromosome number, any phenotypic traits, including those used for characteristic of advanced taxa, could theoretically be studied with our approach, identifying genetic variants for trait determination.

## Results

### Chromosome number statistics

In mammals, the chromosome number changed variously among mammalian species [16], ranging from 2n=102 (*Octomys mimax*) [17,18] to 2n=6 (female in *Muntiacus muntjak*) [19], with the putative number of 2n=46 in mammalian common ancestor [16]. In present work, we collected the chromosome number from 96 mammals (Table S1) ranging from 2n=6 (female in *Muntiacus crinifrons*) to 2n=84 (*Ceratotherium simum simum*).

### Kmer frequencies calculation and quality control

The AF-based software of CVTree [11] was used to construct the phylogenic tree of the 96 mammals. The species was selected on the basis of the genome availability in public database and the chromosome number availability in the previous studies (*Materials and Methods*, Table S1). The set of these 96 species contain the representative of four orders (Cetartiodactyla, Carnivora, Rodentia and Primates). The phylogenic analysis based on AF sequence alignment showed that k=7 oligopeptide (heptapeptide) was able to obtain the best evolutionary tree under four branches (Fig. 1). The human protein sequences were used for heptapeptide-grouping because of the quality of the genome and the completeness of the annotation. According to the quantify control criteria, a total of 82816 groups of heptapeptides which included 2529430 types of unique heptapeptides were picked up for the further analyses.

**Figure 1.**
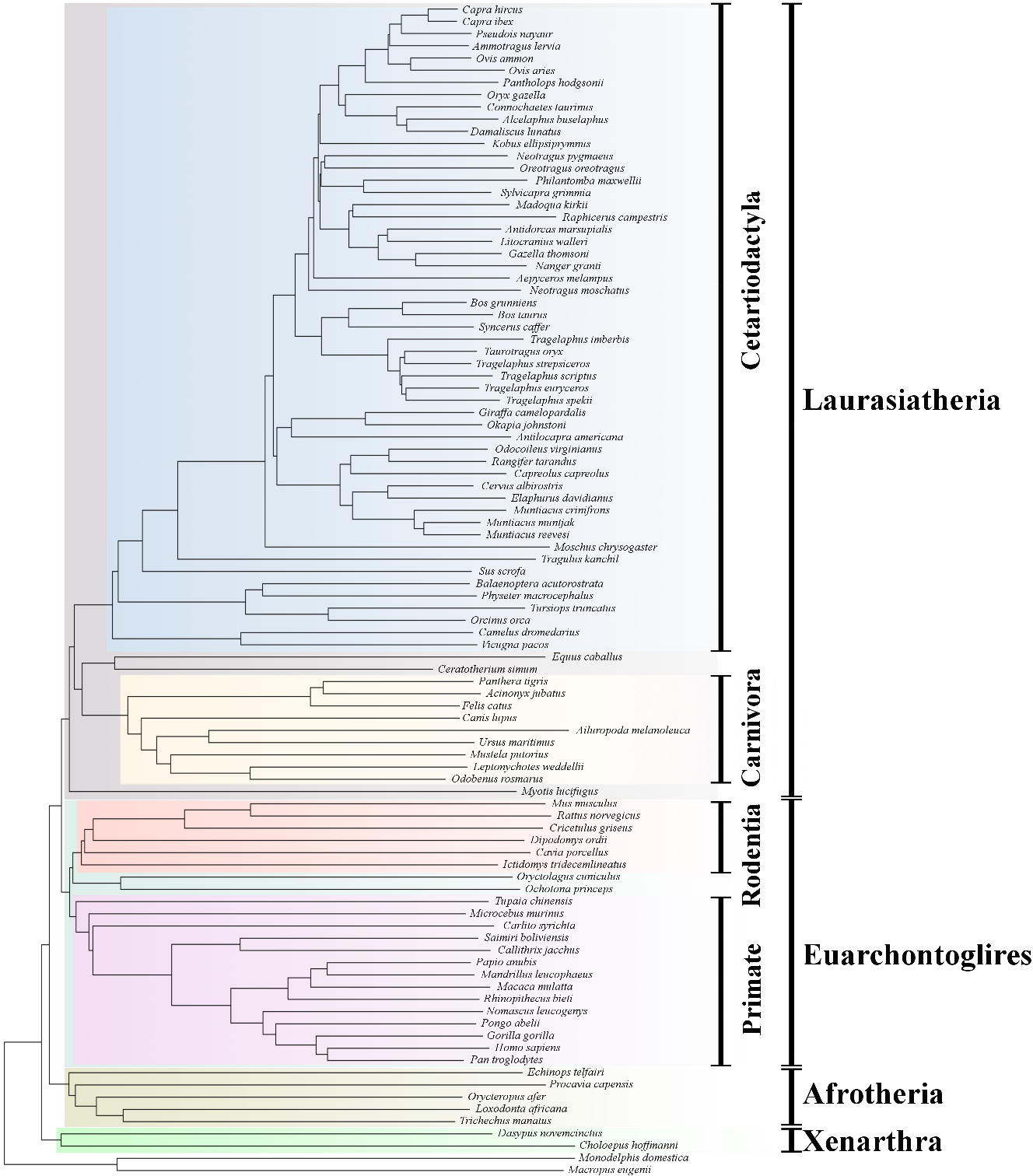
Optimization for the length (k value) of orthologous oligopeptide. The NJ tree is given by the value of k=7. Although the inner topology may be ambiguous or conflict, the 4 orders and the 4 superorders of placental mammals shown with different colors were clustered right. And the relation of the eight lineages were acceptable. The exception for position of superorder Xenarthra could be seen.

#### Assessment of kmer contribution to chromosome number variance

Using random forest method, we estimated the contribution of nearly 2.5 million heptapeptides to the variance of the chromosome number within the mammalian linage. The results indicated that the total heptapeptides explained 18.1% of the chromosome number variance， which verified the substantial role of the heptapeptide copy number for the chromosome number variation.

#### Association analysis

For the association analysis, the Manhattan and QQ plots shown in Fig.2. Table 1 present the significant genes detected by our new approach, including their starting and ending positions in the human genome, the ensemble IDs, the normal-P values and the q values after FDR test. In total, we discovered 6 proteins from 5 genes associated with chromosome number (normal P-values < 0.001 and FDR q-value < 0.05). Of these five genes, the *BAD* had a normal-P value of 1.32E-06, showing the strongest evidence of association with the chromosome number. Moreover, a total of 130 proteins which belong to 84 genes were identified to be associated with the chromosome number (P values < 0.001 without FDR test) (Supplemental Table S2).

**Figure 2.**
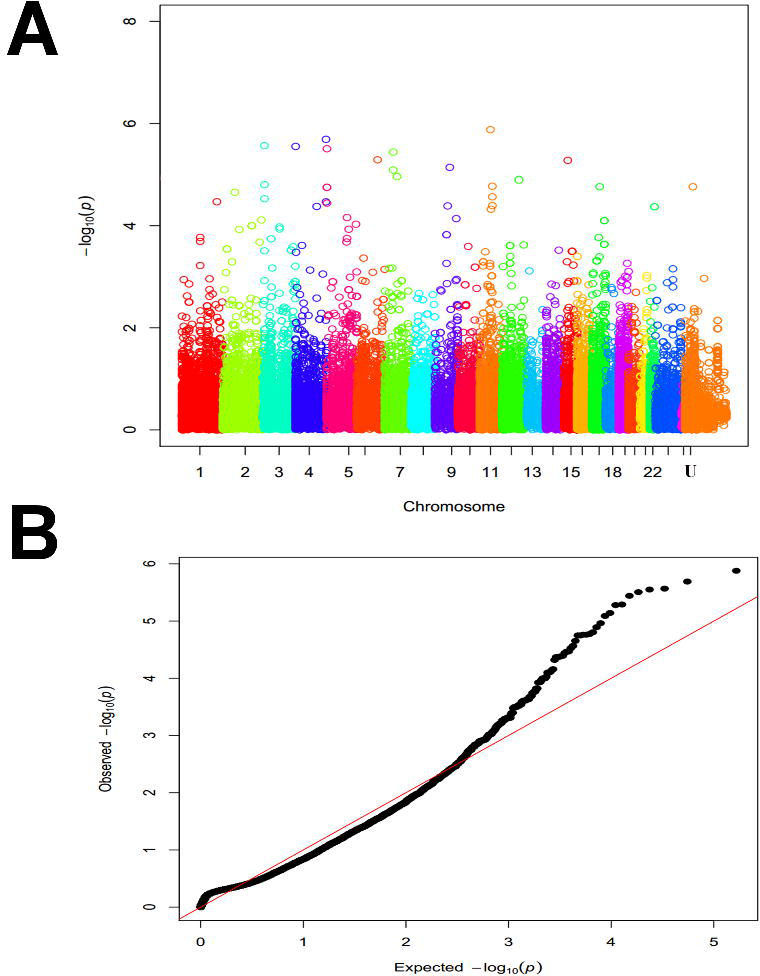
Overall results of the EGWAS. **A**. the plots of Manhattan, the 25 chromosomes (including 22 autosomes, X, Y and the non-chromosome) are color coded. **B**. the Quantile-Quantile plots of −log10 (p values) for chromosome, the observed negative logarithms of the p values in association study are plotted against their expected values under the null hypothesis of no association with the phenotype.

#### Relations between oligopeptide copy number to domain function

Of the 130 significant proteins, the positions covered by the heptapeptides are mainly intrinsic disorder (ID) comparing to other pfam domain regions (Figure 3A). The comprehensive roles of ID in protein interaction network has been discovered in the previous studies [20,21]. Some investigations indicated a crucial role of ID for P53-induced apoptosis [21]. P53 is one of the most well known and intensively-studied tumor suppressors, which plays the central role in apoptosis responding to a broad variety of cell stresses including the DNA double-strand breaks (DSB). Using P53 as an example, we found ID plays a role of bridge between oligopeptides and proteins (Figure 3B, 3C). In Figure 3B both dog and Chinese hamster are species with the chromosome number which is distinct from mammal ancestor, while human chromosome number are quite close to mammal ancestor. It could see that the majority of kmers (except “SGTAKSV”) have a consistent trend of frequencies in dog and Chinese hamster whose chromosome number has deviated from mammal ancestor. There were 4 kmers mapped in the ID region of the gene *P53*, all of which appeared also in ID region of other proteins in human proteome (Figure 3C). These proteins with the same ID region may contribute to the function of P53.

**Figure 3.**
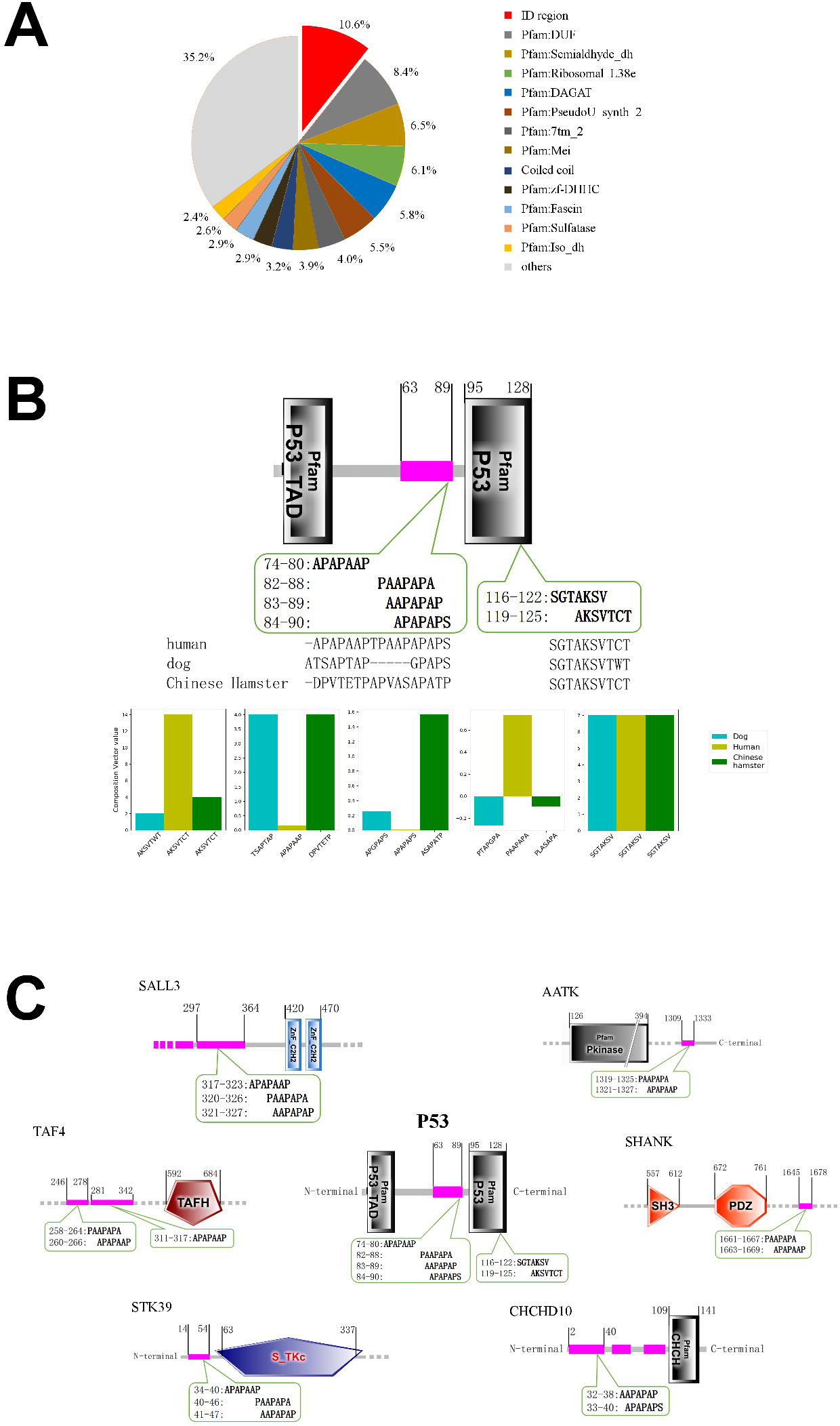
The relation from oligopeptide to function of proteome with the ID as a bridge. **A**. The ratio of ID in all domains mapped with significantly associated heptapeptide. **B**, **C**, And **D** Showed the detailed result of the P53 protein (ENSP00000426252). The coverage of six heptapeptides on the functional domain in human protein are indicated in the center, involving a ID region (purple) and the C-terminal of the P53 PFAM domain. The position of the domains and the oligopeptides in proteins were indicated. The four heptapeptides in the ID region were also appeared in other 72 proteins (supplementary Table S4), six of which covered by at least two of the heptapeptides were shown around. None of the 72 was in the associated gene set, but some of them have had bound evidence to be functional related with P53 such as CDKN1C40.

#### Function enrichment results

To further analyze the function of the significant genes, we conducted the KEGG pathway analysis by using the Genetrial2 platform. The results showed that these 130 proteins were enriched in cancer (hsa05200) and Glycerophospholipid metabolism (hsa00564) (Table S3). The Related genes of *BAD*, *TP53* and *BRCA2* have been proved to play an essential role in cell apoptosis induced by DSB (Figure 4A); and the genes of *GDKQ*, *PLD1* and *CHPT1* are related with the biosynthesis and the metabolism of Phosphatidic acid, which also takes part in the pathways of apoptosis (Figure 4B).

**Figure 4.**
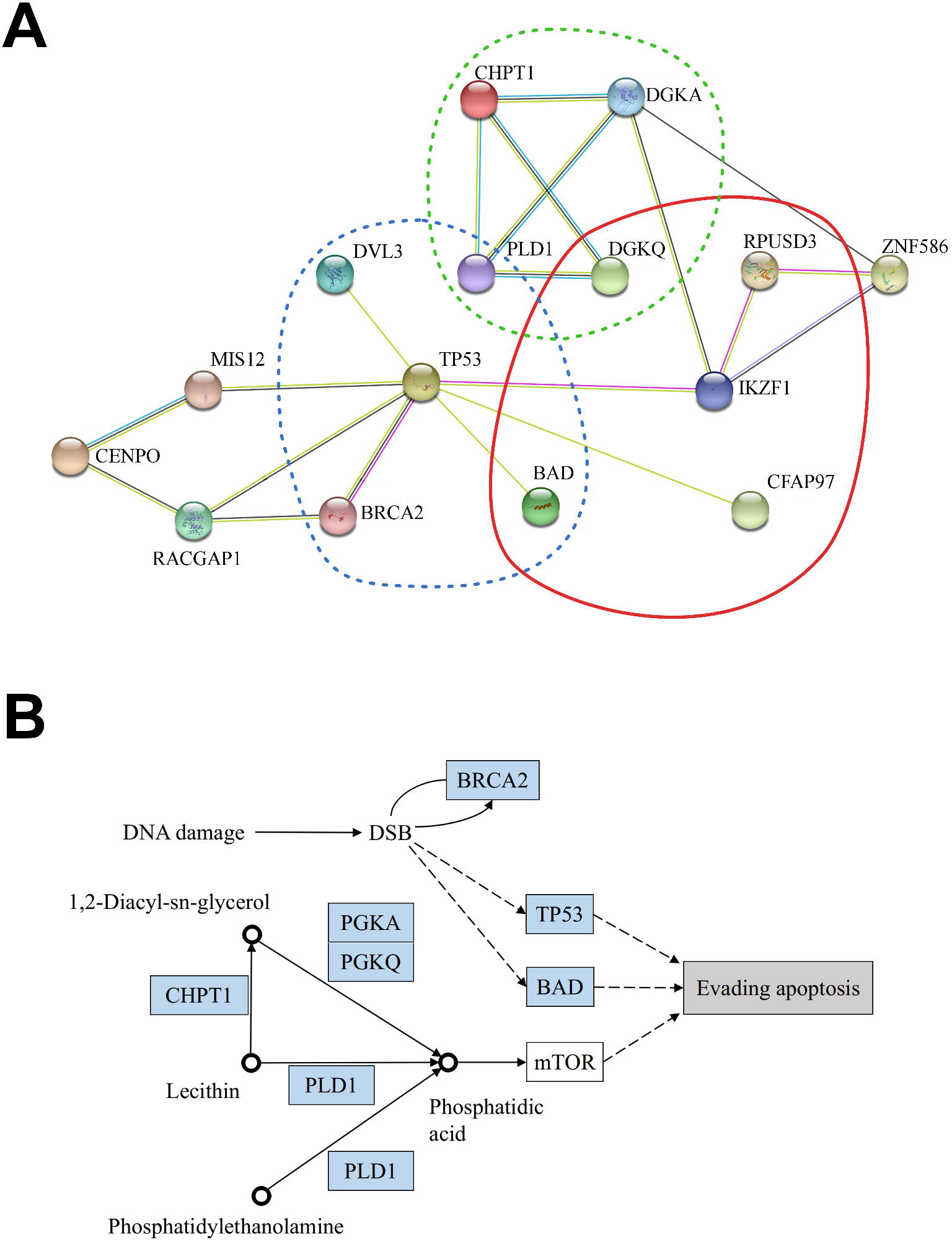
Intensive function analysis for the candidate associated genes. **A**. The refined protein interaction network of related genes modified from STRING website. The balls indicated proteins while the lines connecting balls indicated interaction between proteins. The colors of balls and lines were default lengends from STRING result. The genes in the red solid rings in the right indicated the five genes in the core associated gene set. The region marked with blue dotted lines in the left indicated the five genes in the pathways related to cancer. The region marked with green dotted lines above indicated the four genes in the pathways of Glycerophospholipid metabolism. **B**. The related genes and apoptosis modified from KEGG. The black circle denoted to Glycerophospholipid molecules with names and molecule structures marked nearby respectively. The gene encoding enzymes for related chemical reactions were illustrated in blue boxes. The majority molecule related to apoptosis in cancer was phosphatidate (phosphatidic acid). TP53 and BAD may play essential role during the process to evade apoptosis.

## Discussion

We performed a gene-based GWAS analysis on 96 mammal species in this study. However, a critical constraint in GWAS is the sample size, which is also a potential constraint for the association study in large-scaled inter-species trait. Typically, a GWAS study needs a very large sample size to detect loci varies from phenotype to phenotype, because each SNP makes such a small contribution to the whole variability in phenotype of the trait [15,22]. In the case of large-scaled inter-species trait, the species number should also be large. However, only 96 sequences with chromosome numbers were available in the public databases. With the further development of sequencing technologies, more and more high-quality genome could be acquired. this will further provide more data for our study.

In this study, the kmer frequencies were used to describe the overall feature of genome, which is similar to the AF sequence comparison studies. Generally, the kmer used for AF methods can be either oligonucleotide or oligopeptide so that both genome/transcriptome and proteome data could be used for AF phylogeny study. However, computational complexity was extremely high with oligonucleotide of kmer due to the large size of the mammal genomes. So, many AF phylogeny studies always choose to select oligopeptide in proteome as kmer for complicated multicellular organisms [23]. Here, the oligopeptide frequencies were extracted through CVTree program [7] for association study from the mammalian proteome data. In addition, another essential issue is to determine what length of kmer could reflect the overall feature of proteome. In CVTree, each length of oligopeptide could obtain a distance tree and be filtered out by the criterion based on the accepted phylogenic topologies. As mentioned above, the genetic variation determining the variation of large-scaled inter-species traits was also recorded in kmer characteristics. So, it is reasonable that the length of kmer whose frequencies could obtain the best tree should also obtain the most mount of information of variation to determine the trait. Therefore, we used the length of kmer which determined the best tree for further trait association study.

The connections between the oligopeptide copy number and the targeted proteins, and between the target proteins and the biology context of the trait determination needs further explanation. The heptapeptides are enriched mainly in low complexity or intrinsic disorder (ID) [24]. These unstructured regions in protein have been discovered to be far more important than previous considered [25]. Critical functional involvement includes phosphorylation [26], alternative splicing [27], protein-protein interaction in interactomes [28], and so force. Therefore, the process cat be modeled in three steps. Firstly, the changes of the heptapeptide copy number caused the changes of the ID region appearing in different proteins. Secondly, different ID region distribution in different proteins produces different protein-protein interaction or other function alteration of the target proteins. Finally, the altered function of the related proteins in the proteome caused the variation of the traits.

Further explanation on the chromosome number determination is required, particularly on the function enrichment result. The changes of chromosome number need to be fixed in population, during which the DNA double-strand breaks (DSB) may increase in the cells of individuals in the population. As we know, the increasing DSB may trigger apoptosis. Therefore, for the survive of individuals with different chromosome number, a mechanism which can evade apoptosis under high level of DSB is required. The critical role of DSBs during cancer development [29,30], peculiar the most recent advances in the relation of DSBs and chromothripsis [31], has offered some indirect evidences for identifying the role of DSBs on molecular mechanism about the changes of chromosomes number in evolution.

To better explain the whole-genome role in trait determination, we introduce an analogy involving the comparison of statistical physics. For system consist of large number of microscopic particles, although the motion of single particle could be accurately described with mechanics principle, the macroscopic phenomenon must be quantified with the average statistical feature of particles [32]. As far as the genome case is concerned, it is the similar one. Although the function of single gene obeying to central dogma, the macroscopic heritable trait was an average result of a large number of genes in the complex interaction network. In physics, it has been an implicit view of point that dynamics of single particle could not explain the macroscopic feature of a system with numerous particles [32]. In biology, however, it is a rather radical or revolutionary opinion that traits in individual level must be determined by global feature of the whole genome instead of the local effects of individual genes.

## Conclusion

By using kmer frequencies as the genomic marker to replace SNPs, we extended genome-wide association studies for genetic analysis of the large-scale inter-species trait. Such an approach can be used not only for identifying traditional complex traits within species, but also the large-scaled inter-species complex traits. In this work we took chromosome number variation within mammals as a case to show the availability and the advantages of this approach. We offered a biological explanation for how the overall feature of sequences could participate in the determination of traits, which is the kmers associated with certain phenotype may cover a stretch of continuous polypeptide sequences in protein sequences. These stretch of sequence may work as some functional domain such as ID region which changes its location and the copy number to influence the protein-protein interactions. So that the pattern of the protein interaction network may be changed. And finally the phenotype may be affected. This study has provided a practical method for genetic analysis of the large-scale inter-species trait.

## Materials and methods

### The acquisition and filtration of protein sequence

We selected 96 mammals with annotated proteome data. The newly artiodactyla genome project offered a dataset of 40 newly sequenced genomes (unpublished). Using its annotation system, 8 genome assemblies from NCBI were re-annotated to obtain a refined proteome sequence. 48 proteome sequences were directly downloaded from NCBI.

For kmer frequency analysis, protein sequence from the 96 proteomes were filtered according to two criteria: 1) the length of protein should be longer than 150 amino acid residues, and 2) For multiple proteins from one gene caused by alternative splicing, we only selected the longest one.

### Determination for the length of oligopeptide

The alignment-free methods for evolutionary tree construction were used to determine the best *k* value of the orthologous oligopeptide for copy number counting. Although not applied as widely as alignment method such as BLAST, alignment-free methods have long been developed to analyze sequences [16,33]. Different researchers use different terms to description a class of similar methods. Here we used the well-known term of kmer frequency in computational biology. One disparity is that kmer in computational biology usually denotes oligonucleotides, while in alignment-free method it could denotes either oligopeptide or oligonucleotide. These methods counted the frequencies of kmers for DNA sequence (or protein sequence), defining composition vector with the frequencies to describe the statistical feature of the sequence, constructing a random background to exclude from the vector, and then defining a measurement as the evolutionary distance to construct a distance matrix for concerned sequences (species). Finally NJ tree could be used to describe the evolutionary relation of these sequences.

We used the CVTree to optimize *k* value for the mammalian proteome data above. Considering the restriction of the computer power, CVTree sets the *k* value of protein sequence from 3 to 14. Different *k* values could generate different NJ trees. The random background was defined as a Markovian chain which could be excluded by subtraction parameter. The optimized criterion is designed with three aspects: 1) the four superorders within Eutheria should be correctly clustered, which are Euarchontoglires, Laurasiatheria, Xenarthra and Afrotheria. 2) the monophyletic orders within Euarchontoglires and Laurasiatheria should be correctly clustered, including Carnivora, Cetartiodactyla, Chiroptera and Perissodactyla in Laurasiatheria; and Glires and Primates in Euarchontoglires, and 3) the phylogeny of the monophyletic orders within their respective superorders should be consistent with the general accepted phylogenomic tree. All of the kmer frequency for each species and the distance matrix for NJ tree were the output results by the CVTree program.

### Definition of protein-kmer matrix and quality control

In GWAS, it is a general strategy to group SNP within one gene (or transcript) together to perform linear regression. Here we used the similar methods with the aim of grouping oligopeptides into protein sequences to identify the association between protein and the phenotype. We used the protein sequences of the unfiltered human proteome downloaded from the Ensembl Genes version 90 as the reference to group oligopeptides. Each group corresponds to the oligopeptides appearing in one human protein sequence. Theoretically, the grouping should be random. But this would produce a vast number of groups which is impossible for association computation.

In CVTree results, the frequency value of −1.0 or 0.0 was used to denote missing data. We removed these data for oligopeptide groups with the following criterion: 1) All kmers with the missing data < 90% and all transcripts with kmers <5 were filtered out, 2) All the species with the missing data >10% were ruled out for each transcript.

### Phenotypic preparation

The mammalian karyotype has been extensively investigated since 1960s [16]. Since ever, some species were renamed, or their taxonomic places were changed. This makes some difficulties to get the chromosome number information of the species with the available proteome data. We manually checked related literature for these data. There are also some species with intra-species variation of the chromosome number, the numbers are rounded to the nearest integer average for these cases. We can see the details of chromosome number in Supplementary Table S1. To eliminate the influence of species difference, the phenotypic values of the chromosome number was adjusted by the evolutionary relation of the 96 mammals. In other words, the residuals after fitting the above distance matrix from CVTree were treated as the phenotypic values of traits for the further association studies.

### Variance component analysis

The analysis is conducted through the R script programmed with R languange version 3.4.3. As the core part of the program, RandomForest (version 4.6-14) was used for the percentage of variance explained calculated which can be directly provided from the R script outcome.

### Association analysis

For each group (protein of transcript) of oligopeptides, we constructed principal components (PCs) according to the following rules [14]. We treated each oligopeptide (kmer) within a group as a single variable, and calculated the variance-covariance matrix. Then we calculated the eigenvalues and the eigenvectors of the covariance matrix. PCs were obtained by multiplying the eigenvectors by the kmer frequency matrix. Thus, PCs were mutually independent within a protein sequence, and could be treated as the independent variables for regression analysis and significance tests. After these steps, multiple regression analysis for the adjusted chromosome number on the oligopeptides (kmers) was turned to a simple regression by using the PC. A simple correlation coefficient corrected by chromosome numbers and PCs could be used to calculate the P-value of the PC.

In the Fisher’s combination test, χ^2^ was constructed by combining K independent p values for PCs, as follows:

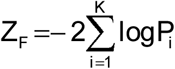

This test statistic followed a χ^2^ distribution with the degree of freedom 2ↃK under the null model, and a new p-value was calculated from this Chi-square distribution for each transcript.

The false discovery rate test was adopted for multiple tests of all groups (proteins) in the proteome. Proteins whose nominal q value less than 0.05 was considered as core associated gene set at the proteome-wide significance level. Besides, proteins with nominal p value less than 0.001 was categorized under the peripheral associated gene set.

### Case study of ID region in P53

In the association study, one of P53 transcripts (Ensembl number ENSP00000426252) was used and showing significant association with chromosome number, in which 6 heptapeptide were selected after filtering absences in the 96 mammals. So, the selected three species with the representative chromosome number, human (2n=46), dog (2n=78), Chinese hamster (2n=22) showed the subtracted heptapeptide frequencies. The heptapeptide locations were identified in SMART (URL see below). The frequency was subtracted with the random background in CVTree, and the result values were compared among the three species. Not that the heptapeptide of “AAPAPAP” was absent in dog and Chinese hamster, so only five were shown in Figure 3B. For all other proteins, the domain identification of proteins was all performed in SMART.

### Function enrichment and analysis

We used the peripheral associated gene set for a function enrichment with Genetrial2 and for a protein interaction network analysis with STRING. For Genetrial2 analysis, the enrichment algorithm used was Over-representation analysis, the adjustment method was FDR adjustment offered by Genetrail2 with significance level of 0.1. We excluded results of pathways which include no gene from the core associated gene set. For STRING analysis, the low confidence for minimum required interaction score (>0.150) was used. The subcellular location information was obtained in GeneCards.

### Urls used in Methods

NCBI, https://www.ncbi.nlm.nih.gov/assembly. Ensembl http://www.ensembl.org/index.html.

SMART, https://smart.embl-heidelberg.de/. Genetrial2, https://genetrail2.bioinf.uni-sb.de/.

STRING, https://string-db.org/. GeneCards, http://www.genecards.org/.

### Authors’ contributions

Qi Wu and Fuwen Wei designed the research. Qi Wu and Lei Chen prepared the data. Qi Wu and Huizhong Fan performed the research and interpreted the data. Qi Wu, Huizhong Fan, Yibo Hu and Fuwen Wei wrote the manuscript.

### Competing interests

The authors declared no competing interests of this work.

## Acknowledgments

We thank Prof. Qiang Qiu for the helpful discussion of the project and Dr. Xinhai Li for the help and assistance in random forest method application. This work is supported by grants from National Key Program of Research and Development, Ministry of Science and Technology Grant (No. 2016YFC0503200), the Strategic Priority Research Program of the Chinese Academy of Sciences (No. XDPB0202), the Key Research Program of the Chinese Academy of Sciences (No. KJZD-EW-L07) and the NSFC Major Scientific Research Program (No. 91746119).

